# An RNA-based system to study hepatitis B virus replication and select drug-resistance mutations

**DOI:** 10.1101/787630

**Authors:** Y. Yu, W.M. Schneider, E. Michailidis, A. Acevedo, Y. Ni, P. Ambrose, C. Zou, M. Kabbani, C. Quirk, C. Jahan, X. Wu, S. Urban, A. Shlomai, Y.P. de Jong, C.M. Rice

## Abstract

Hepatitis B virus (HBV) chronically infects over 250 million people worldwide, increasing their risk of liver cirrhosis and hepatocellular carcinoma. There is a vaccine to prevent new infections, but no efficient cure for chronic infection. New insights into HBV biology are needed to improve cure rates for this widespread devastating disease. We describe a method to initiate replication of HBV, a DNA virus, using synthetic RNA. This approach has several advantages over existing systems: it eliminates contaminating background signal from input virus or plasmid DNA and can be easily adapted to multiple genotypes and mutants. Further, it can be applied to identify anti-HBV compounds, measure anti-HBV drug efficiency, study virus evolution, and, as we demonstrate, it can be uniquely applied to predict antiviral drug resistance.

## Main Text

HBV is a small DNA virus, transmitted by blood or other bodily fluids, which infects human hepatocytes. Upon HBV entry into hepatocytes, the relaxed circular, partially double-stranded DNA (rcDNA) genome is converted in the nucleus to the stable episomal form known as covalently closed circular DNA (cccDNA) (Fig. S1A). cccDNA is transcribed by host RNA polymerase II (Pol II) to produce all HBV RNAs (Fig. S1B), and cccDNA persistence in hepatocytes is thought to be responsible for chronicity (*1, 2*).

Currently, chronic HBV-related disease claims an estimated 880,000 lives each year (*3*). There is an effective vaccine to prevent new HBV infections, but it does not benefit those already infected and vertical transmission remains a problem in some areas of the world. Standard therapies for chronic HBV include injectable interferon alpha (IFNα)—a naturally occurring cytokine that elicits a broad antiviral response—and orally administered nucleoside/nucleotide analogs. IFNα therapy can cure ∼10% of patients, but it elicits often intolerable side effects and since few patients benefit, it is seldom used. In contrast, nucleoside/nucleotide analogs that effectively suppress HBV replication are well tolerated, but they do not eliminate the virus and therefore therapy is indefinite and often lifelong. New therapies are critically needed to improve HBV cure rates.

Lately, there has been renewed interest in curing chronic HBV. This has been driven, in part, by recent success in curing chronic hepatitis C virus infection and by the discovery of the HBV entry receptor, NTCP (*4, 5*), which has made it possible to study the full HBV lifecycle in cell culture. As a result, there is a growing belief that with new anti-HBV therapies and armed with a better understanding of HBV biology, curing chronic HBV may be within reach.

But despite progress, existing cell culture-based methods to study HBV have several limitations. For one, infection *in vitro* requires a high virus particle to cell ratio which results in a high background of contaminating viral DNA and protein from the inoculum. Background contamination also plagues methods that initiate HBV replication with either plasmid DNA transfection or using cell clones with integrated HBV genomes. In these cases, much of the viral RNA and protein originate not from the authentic viral template, cccDNA, but from either the transfected plasmid or integrated DNA. In fact, expression of viral RNAs and proteins is high even when the HBV genome encodes catalytically inactive HBV polymerase (Fig.S2).

To address these challenges, we developed a method to initiate HBV replication with *in vitro*-transcribed RNA. The HBV genome is reverse transcribed from an RNA template— pregenomic RNA (pgRNA)—and we reasoned that initiating replication and gene expression with RNA would have several benefits over systems that initiate replication with plasmid or integrated DNA. Initiating replication with RNA would, for example, eliminate contaminating HBV DNA that confounds qPCR reactions, and since not all viral RNAs are required to initiate replication, some viral proteins (e.g. HBsAg, a commonly used marker for infection) would be produced only if the viral lifecycle progressed and cccDNA was established.

We chose to launch HBV replication with pgRNA alone since pgRNA is not only the template for reverse transcription, but also the template for translation of the HBV core and polymerase (Pol) proteins, both of which are required for reverse transcription (Fig. S3). The idea to initiate HBV with pgRNA is supported by previous work on a relative of HBV, duck hepatitis B virus (DHBV), with the observation that DHBV pgRNA can initiate infection in cultured cells (*6*).

**Figure 1** demonstrates that transfected HBV pgRNA is translated to produce the core protein (HBcAg), reverse transcribed by the viral polymerase (Pol), and transported to the nucleus to form cccDNA, yielding HBsAg expression. Southern blot on Hirt-extracted DNA isolated from pgRNA-transfected cells confirms that pgRNA gives rise to cccDNA. Transfection of pgRNA encoding a catalytically inactive Pol mutant (YMHD) demonstrates that the HBV DNA and HBsAg signal is replication dependent and that there is minimal to no contaminating background signal from input RNA.

**Fig. 1.**
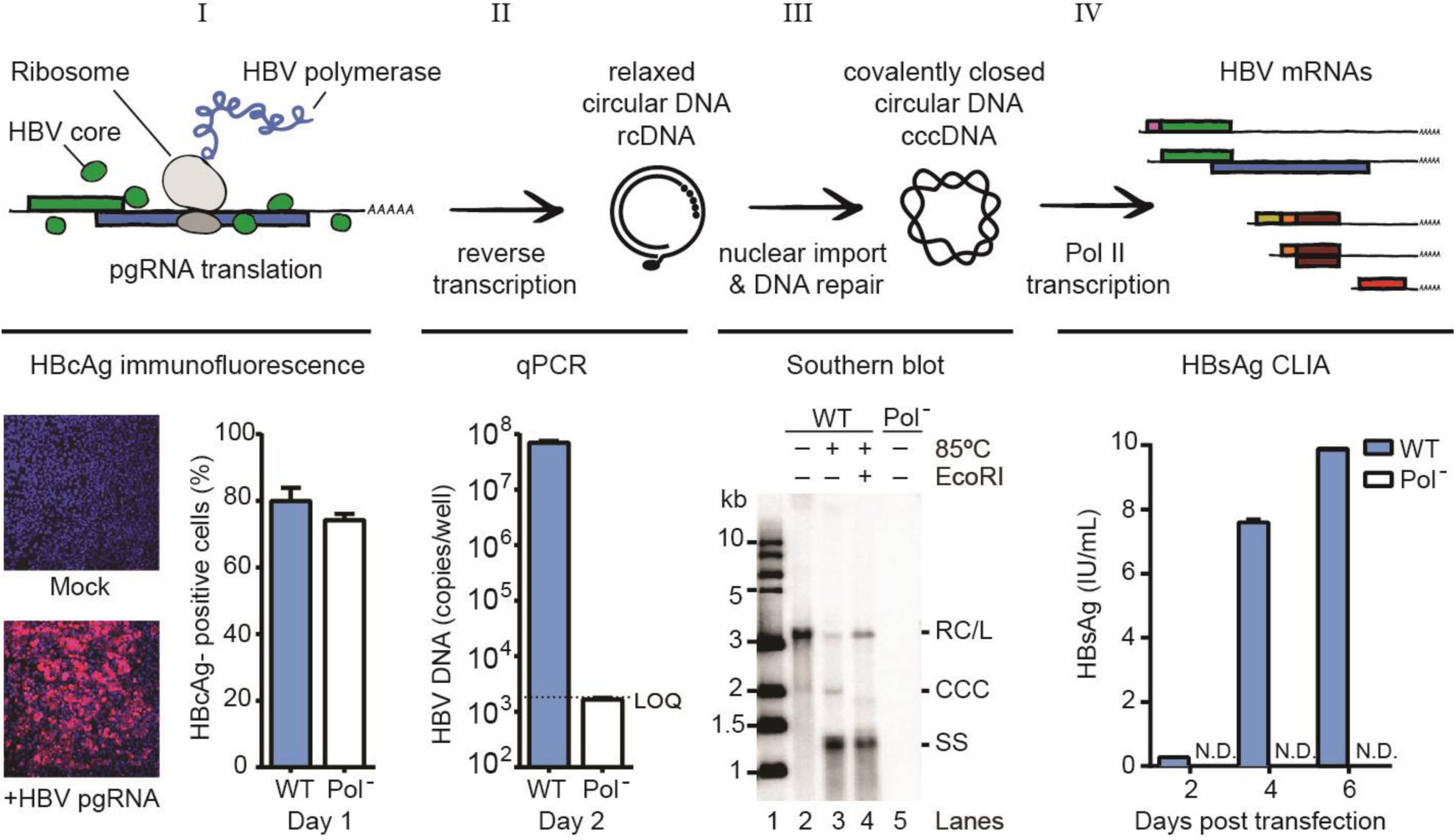
The RNA launch system. From left to right, the figures shows that *in vitro*-transcribed pgRNA is (I) translated (HBcAg is detected by immunofluorescence in ∼80% of Huh7.5-NTCP cells), (II) reverse transcribed (qPCR detects HBV DNA and the signal is HBV polymerase-dependent, as a catalytic site mutation (Pol^--^, YMHD) decreases the DNA signal by 10,000-fold), (III) is transported to the nucleus to form cccDNA (Southern blot confirms cccDNA is produced; see below for lane description), and (IV) cccDNA is transcribed to produce viral mRNA and protein (shown by HBsAg chemiluminescent immunoassay (CLIA)). HBcAg, HBV core antigen; HBsAg, HBV S antigen; WT, wildtype; LOQ, limit of quantification; IU, international units; N.D., not detected; copies/well, one well of 6-well plate; kb, kilobase; Southern blot: Lane 1, 1-kb ladder; Lane 2, WT HBV DNA; Lane 3, WT HBV DNA heated to 85°C; Lane 4, WT HBV DNA heated to 85°C then linearized with EcoRI; Lane 5, Pol^--^HBV DNA; RC/L, relaxed circular/linear DNA; CCC, covalently closed circular DNA; SS, single stranded DNA.

Until recently, most HBV cell culture studies have focused on genotypes A and D. Similarly, our initial work was performed with genotype A HBV. There are, however, additional HBV genotypes, several of which are highly prevalent in various parts of the world (**Fig. 2A** and table S1) (*7*). It is increasingly recognized that studying multiple genotypes is important, since not all HBV genotypes respond equally well to anti-HBV therapies (*8*). We next tested whether the RNA launch method could be used to study multiple HBV genotypes. For this, we cloned representative pgRNA sequences for HBV genotypes B-H downstream of a T7 promoter and *in vitro*-transcribed pgRNAs. We found that transfected pgRNA from all eight HBV genotypes yielded HBV DNA and all but genotype G yielded HBsAg (**Fig. 2B-C**). As expected, the signal is replication dependent since the reverse transcriptase inhibitor, entecavir (ETV), decreases HBV DNA levels and eliminates HBsAg production. This demonstrates that the RNA launch method can be applied to study multiple HBV genotypes and viral mutants without the high background signal typical of plasmid transfection and virus infection approaches. Further, this approach could be used to bypass virus entry and study how genotype differences affect post-entry steps in the virus lifecycle. For example, genome amplification, accumulation of replication intermediates, and cccDNA formation. Indeed, the results in **Fig. 2** and S4) indicate there are differences in reverse transcription and HBsAg production among genotypes that can be further explored with this system.

**Fig. 2.**
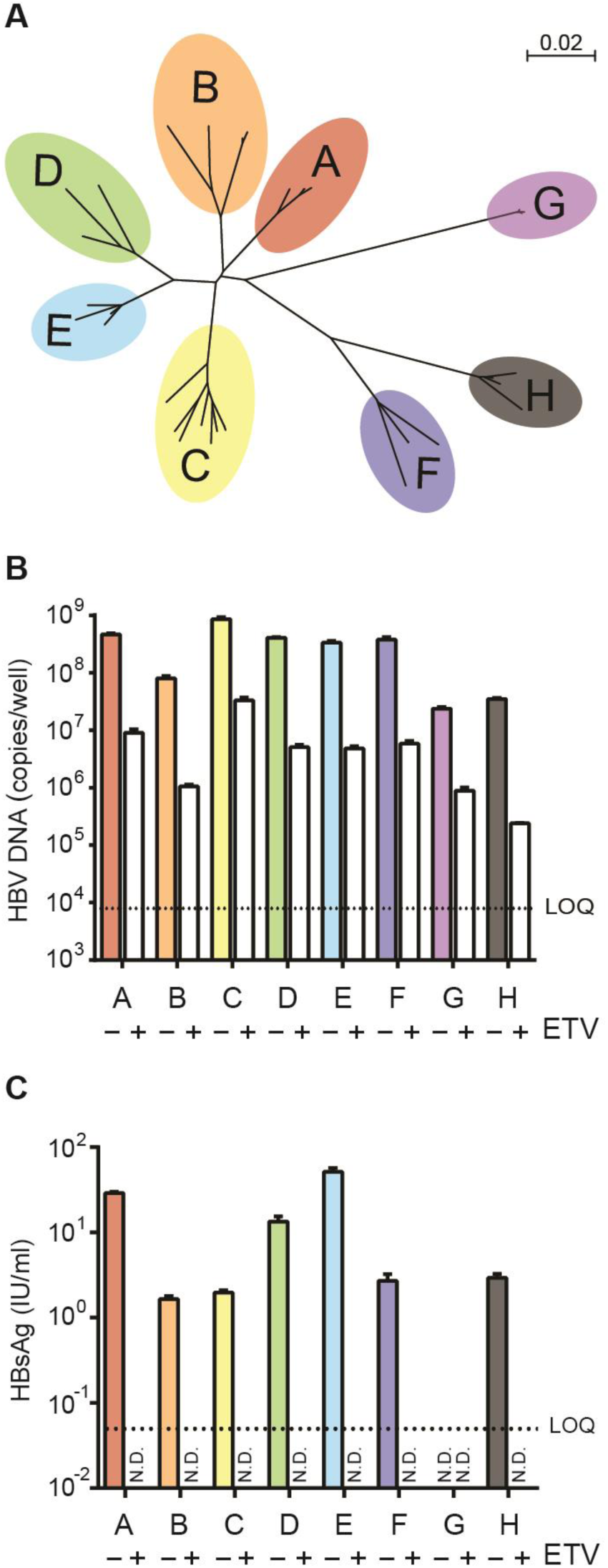
Pan-genotype HBV replication from *in vitro*-transcribed HBV pgRNA. (**A**) Unrooted phylogenetic tree of HBV genotypes A-H. Tree was generated using SeaView (*22*). Scale bar indicates nucleotide substitutions per site. (**B**) qPCR of intracellular HBV DNA for HBV genotypes A-H two days post transfection with and without 10 μM ETV. (**C**) CLIA detects secreted HBsAg from HBV genotypes A-H six days post transfection with and without 10 μM ETV. ETV, entecavir; copies/well (6-well plate); N.D., not detected; LOQ, limit of quantification.

We next tested whether the RNA launch method could be used to compare the potency and effectiveness of anti-HBV inhibitors. For this, we compared three reverse transcriptase inhibitors known to inhibit HBV: ETV, tenofovir disoproxil fumarate (TDF), and lamivudine (LAM). As shown in **Fig. 3A** and Table S2, all three drugs inhibited HBV replication in a dose-dependent manner, demonstrating that the RNA launch method can be used to study the efficacy of anti-HBV compounds in a simple and easy-to-use assay. These results also suggest that the method could be used as a screening tool to discover novel HBV inhibitors.

**Fig. 3.**
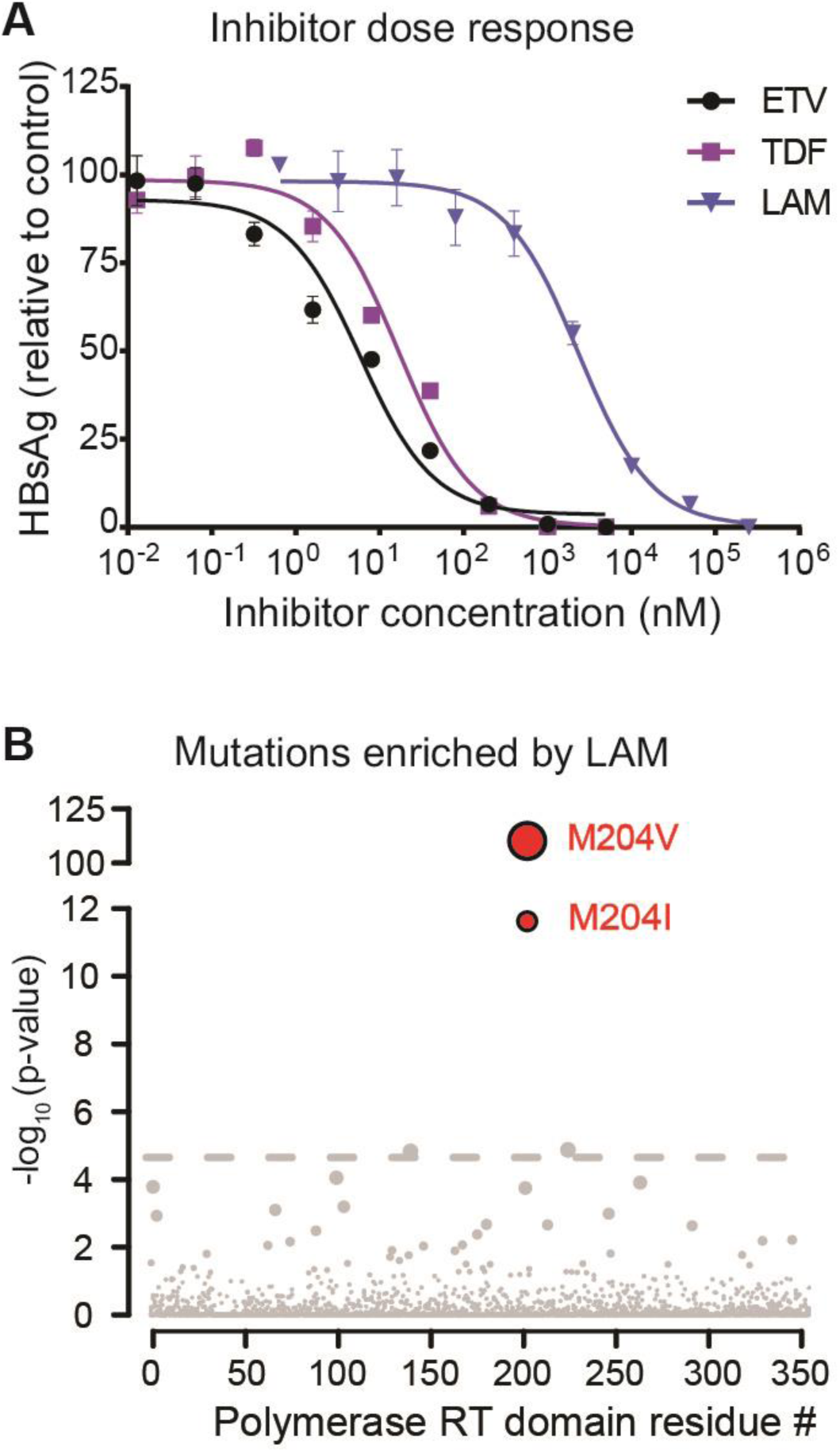
RNA launch method can be used to test drug efficiency and identify drug-resistance mutations. (**A**) HBV reverse transcriptase inhibitor dose response six days post transfection with HBV genotype A pgRNA. Secreted HBsAg was quantified by CLIA and normalized to untreated controls. Data plotted are n=3 (**B**) Sequencing extracellular HBV DNA from cells transfected with HBV pgRNA in the presence of 400 μM LAM identifies the two most common amino acid substitutions, M204V and M204I, found in LAM-resistant patients. Threshold = Bonferroni correction, alpha = 0.05.

Given the low background from input pgRNA, we next asked whether we could use the system to select for drug-resistance mutations. Identifying drug-resistance mutations can provide drug developers with useful information for prioritizing or down selecting compounds and compound series for further development. Moreover, it can provide insight into drug-protein interactions and be used as an iterative approach with new chemistry to improve the resistance profile of a compound series. However, selecting drug resistance mutations for HBV is challenging since the virus spreads too poorly *in vitro* for drug-resistant variants to emerge. Consequently, there is no cell culture-based method to assess antiviral resistance for anti-HBV compounds; however, with new anti-HBV drugs currently undergoing preclinical and clinical evaluation, there is a high demand for methods to assess drug resistance.

We reasoned that several features of the RNA launch method would enable us to enrich HBV drug-resistant variants. For one, unlike protocols that initiate infection with plasmid DNA, sequence diversity is inherent in the pgRNA population. This is because T7 polymerase, which we use to transcribe HBV pgRNA, has an error rate of ∼10^−4^ (Fig. S5B) and therefore produces a diverse mixture of sequence variants immediately available for selection (*9*). Also, since transfecting pgRNA bypasses the entry step, cells likely sample more genomes than they would with other HBV systems that initiate infection with virus particles. And further, since drug-resistant genomes can be enriched by PCR without contamination from input plasmid or viral DNA, we reasoned it would be possible to identify mutations that confer drug resistance by deep sequencing. This concept is outlined in Fig. S5A.

As a proof-of-concept for this approach, we tested whether we could enrich for LAM resistance mutations from *in vitro-*transcribed pgRNA. Two days after transfecting Huh-7.5-NTCP cells in triplicate with pgRNA we harvested virus-containing cell supernatants and isolated viral DNA. Viral DNA was then amplified by PCR (Fig. S4C) and sequenced by MiSeq. Remarkably, we found that with only a single round of selection, the two most common clinically-relevant LAM-resistance mutations, M204V and M204I, were enriched in the population (**Fig. 3B**) (*10*). This demonstrates that the RNA launch method can be used to identify mutations that confer drug resistance. For antivirals with high resistance barriers this approach can be modified to include mutagenized templates and mutant polymerases with higher error rates to increase sequence diversity. This approach can also be combined with deep mutational analysis and applied to study viral evolution under various selective pressures beyond anti-HBV drugs.

Altogether, the RNA launch method is a unique and versatile new tool to study basic HBV biology, identify novel anti-HBV inhibitors, and inform drug development.

## Acknowledgements

We thank Dr. Haitao Guo (Indiana University) for providing the HepDE19 cell line and Yosef Shaul for the 1.3x HBV plasmid. We thank Kuanhui Xiang, Jessica Sonnabend, Alex Kaufman, Ankit Bhatta, and Jonathan Pabon for technical assistance and Hans-Heinrich Hoffmann for helpful discussion. We also thank the Rockefeller University High-Throughput and Spectroscopy, Genomics, and Flow Cytometry Resource Centers.

## Author contributions

Conceptualization: Y. Y., W.M.S., A.S. and C.M.R.; Formal analysis and Software: A.A.; Investigation: Y.Y., W.M.S., E.M., A.A., P.A., C.Z., M.K., C.Q., C.J., X.W., and Y.P.D.; Resources: Y.N. and S.U. (for providing HBV genotypes B, C, E-H); Supervision: Y.P.D. and C.M.R.; Visualization: Y.Y. and W.M.S.; Writing - original draft: W.M.S.; Writing – review & editing: Y.Y., W.M.S., E.M., A.A., Y.N., P.A., S.U., A.S., and C.M.R.

## Data and materials availability

All data is available in the main text or the supplementary materials.

## Supplementary Materials

### 1. Materials

#### Plasmids

All plasmid sequences used in this study have been uploaded to GenBank (table S1). Briefly, HBV pgRNA sequences were cloned into the pGEM-3Z plasmid backbone immediately downstream of a T7 promoter. The genotype A sequence has been previously described as serotype adw2 (*11*). Genotype D was derived from the HepDE19 cell line (*12*). Genotypes B, C, and E-H were derived from over-length HBV constructs originally cloned by Stephan Schäfer (*13*) and then subcloned into the pGEM3Z plasmid backbone. The HBV-YMHD Polymerase mutant was made in the genotype A backbone.

#### Cells

Huh-7.5-NTCP cells were made as previous described for HepG2-NTCP (*14*) and were maintained in Dulbecco’s Modified Eagle Medium (DMEM, Fisher Scientific, cat. #11995065) supplemented with 0.1 mM nonessential amino acids (NEAA, Fisher Scientific, cat. #11140076) and 10% hyclone fetal bovine serum (FBS, HyClone Laboratories, Lot. #AUJ35777). HepDE19 cells were maintained on collagen-coated plates in DMEM supplemented with 3% FBS, 0.1 mM NEAA, and 2 μg/ml tetracycline.

### 2. Methods

#### Synthesis of HBV pgRNA

10 µg of plasmid DNA was linearized by digestion with NotI-HF. Linearized DNA was purified with the MinElute PCR purification kit (Qiagen, cat. #28004) according to the manufacturer’s instructions, diluted to 0.5 µg/µl in EB buffer, and 2 µg was used as template for *in vitro* transcription. Transcription was performed using the T7 mScript™ Standard mRNA Production System (CELLSCRIPT, cat. #C-MSC100625). The reaction was incubated at 37°C for 75 min followed by 15 min DNase treatment at 37°C. RNA was immediately purified using the RNeasy mini kit (Qiagen, cat. #74014) following the manufacturer’s instructions, including the optional on-column DNase digestion step (Qiagen, cat. #79254). Capping and polyadenylation was performed following the Cap 1 mRNA protocol described in the user’s manual and the RNA was again purified using the RNeasy mini kit without on-column DNase digestion. To remove the residual DNA contamination, 50-60 µg of RNA was further digested by Turbo DNase kit (Fisher Scientific, cat. #AM2238) in 50 µl reaction volume with 5 µl 10X buffer, 3 µl Turbo DNase and 1 µl SUPERase-In (Fisher Scientific, cat. #AM2696). RNA was again purified using the RNeasy mini kit. The RNA yield was ∼45-60 µg per reaction.

#### RNA transfection

Huh-7.5-NTCP cells were seeded at 2.5 × 10^5^ cells per well in 6 well plates two days before transfection. The media was changed to 2 ml DMEM containing 1.5% FBS and 0.1 mM NEAA just before transfection. For each transfected well, 1 µg HBV pgRNA was mixed with 5 µl Lipofectamine™2000 (Fisher Scientific, cat. #11668019) in 500 µl Opti-MEM Reduced Serum Medium (Fisher Scientific, cat. #51985034) and incubated at room temperature for 20 min. The mixture was then added to cells and spinoculated by centrifugation at 1000 x g for 30 min at 37°C. Six hours later the media was removed and replaced with DMEM containing 10% FBS and 0.1 mM NEAA.

#### DNA transfection

To detect HBsAg produced by plasmid transfection from genotypes A-H (table S1), Huh7.5- NTCP cells were seeded at 2.5 × 10^5^ cells per well in 6-well plates and transfected two days later in DMEM containing 1.5% FBS and 0.1 mM NEAA with 2.5 µg of plasmid DNA using X-tremeGENE™ 9 DNA transfection reagent (Sigma cat. #6365779001) at 5:1 reagent:DNA ratio. Supernatants were collected four days post transfection for HBsAg CLIA. To compare features of the RNA launch to plasmid transfection (Fig. S2), a 1.3xHBV plasmid (genotype A) (*15*) was transfected into HepG2-NTCP cells. Cells were seeded at 7 × 10^4^ cell/well in collagen-coated 24- well plates and transfected with 0.5 µg DNA the following day using Lipofectamine™2000 at 3:1 reagent:DNA ratio. Two days post transfection, cells and supernatants were collected for analysis. To compare features of the RNA launch to HepDE19 cells (Fig. S2), HepDE19 cells were seeded at 7 × 10^4^ cell/well in collagen-coated 24-well plates in media containing 2 µg/ml tetracycline. The following day media was changed and replaced with media with and without tetracycline. Five days later supernatants and cells were collected for analysis.

#### Analysis of HBV translation and replication

HBsAg was quantified using a chemiluminescence immunoassay (CLIA) kit (Autobio Diagnostics Co., Zhengzhou, China) according to the manufacturer’s instructions. HBsAg experiments were performed in 24-well format. To quantify HBV DNA, total DNA was extracted from individual wells of 6-well plates using QIAamp DNA Blood Mini Kit (Qiagen, cat. #51106) and HBV DNA was detected by qPCR as previously described (*14*). In addition, we designed a separate set of primers targeting the HBV Core region of genotype A that we used in SYBR green assays. Both assays gave similar results; however, the SYBR assay more consistently yielded a background signal (YMHD polymerase mutant) below the limit of quantification. Primers and probe sequences for qPCR are listed in table S3. For immunofluorescence analysis, cells were stained with Anti-HBc (Austral Biologicals, cat. #HBP- 023-9) at 1:500 dilution or Anti-HBs (Abcam, cat# ab9193) at 1:500 dilution. Fluorescent images were quantified in ImageJ (NIH, Bethesda, Maryland) using the thresholding method, similar to what has been previously described (*16*). Southern blot was performed combining two protocols previously described (*15, 17*) with the several modifications. HBV cccDNA was enriched by Hirt extraction as previously described (*17*). The extract was digested at 37°C for 2 h in 400 µl 1X Cutsmart buffer (NEB) with 40 µg RNase A (Thermo Scientific, cat. # EN0531) to remove RNA and 60 U of HindIII-HF (NEB, cat. # B7204S) to digest genomic DNA. The reaction was stopped by adding 200 µg of proteinase K (Fisher Scientific, cat. #25530015) for 30 min at 37°C followed by phenol/chloroform extraction. DNA was precipitated with ethanol and dissolved in 25 µl TE (10 mM Tris, EDTA 1 mM, pH 8). DNA extracted two, four, and six days after transfection was combined and split into three separate tubes for further treatment with heat and/or EcoRI digestion. Samples were run overnight at 4°C in a 1.2% agarose TAE gel then transferred to a Hybond-XL membrane (Fisher Scientific, cat. #RPN303N) for 36-48 h. ^32^P-labeled hybridization probes were prepared with the Prime-It II Random Primer Labeling Kit (Agilent Technologies, cat. #300385) using linearized 2X HBV DNA as a template as previously described (*15*). Probes were hybridized overnight at 65°C. The membrane was washed as described (*17*) and HBV DNA was visualized by phosphorimager.

#### CirSeq analysis of *in vitro*-transcribed pgRNA

CirSeq of *in vitro*-transcribed pgRNA of HBV genotype A was performed as previously described (*18*). Briefly, fragmented pgRNA was circularized and converted to DNA by rolling-circle reverse transcription, yielding tandemly repeated cDNAs. Two replicates were prepared and 325-cycle single-end sequencing of both replicates was performed on an Illumina MiSeq at the Rockefeller University Genomics Resource Center. Tandem repeat reads were converted to consensus sequences, filtering out random errors generated during library preparation and sequencing, and then mapped to the reference. The error rate of T7 polymerase was determined as the frequency of mismatches with respect to the reference sequence for bases with average quality score >= 20. Sequences are deposited in the Sequence Read Archive, accession number XXX.

#### HBV inhibitor treatment

Entecavir was purchased from Cayman Chemical Company (cat. #209216-23-9), lamivudine was purchased from Sigma (cat. #L1295), and tenofovir disoproxil fumarate was obtained through the AIDS Reagent Program, Division of AIDS, NIAID, NIH. Huh-7.5-NTCP cells were seeded at 1 × 10^4^ cells per well in 96-well plates two days before transfection. One day before transfection, media was changed and replaced with 100 µl DMEM containing 10% FBS and 0.1 mM NEAA with HBV inhibitors or vehicle control. The following day, each well was transfected as described above with 34 ng of HBV pgRNA in media containing drug. Six hours later, media was changed and replaced with drug-containing media. Every two days, media was collected and replaced and 50 µl of collected supernatants were used for HBsAg CLIA, as described above.

#### Enriching LAM-resistant viral variants

To enrich mutations that confer LAM resistance, 2.5 × 10^5^ Huh-7.5-NTCP cells were seeded two days before pgRNA transfection as described above. The cells were treated with 400 µM LAM 24 h before RNA transfection. Transfection was performed and cells were maintained in the presence of 400 µM LAM. Supernatant was collected two days after transfection and concentrated with Amicon-Ultra-0.5 ml 100 kDa centrifugal filters (Millipore, cat. #UFC510096) to a volume of 200 µl. HBV DNA was then extracted using the QIAamp MinElute Virus Spin Kit (Qiagen, cat. #57704). DNA was eluted with 50 µl buffer AVE and amplified by two PCR reactions prior to sequencing. The PCR products from both rounds were separated in a 1% TAE agarose gel and purified by gel extraction (Qiagen, cat. #28704). The first PCR used primers RU-O-26880 and RU-O-24242 to amplify near full length HBV sequence in 100 µl volume using 50 µL 2X KOD start Master Mix (EMD, cat. #71843), 20 µl of eluted DNA, and 15 pmol of each primer. The PCR was perform by (i) denaturation at 95 °C for 1 min (one cycle); (ii) PCR at 95 °C for 15 s, 62°C for 30 s, and 68 °C for 60 s (25 cycles for no drug treatment and 32 cycles for 40 µM and 400 µM LAM treatment) (iii) 68 °C for 2 min and 10 °C to hold. The PCR products were separated in a 1% TAE agarose gel and purified with the MinElute Gel Extraction kit (Qiagen, cat. #28604) and eluted in 15 µl EB buffer. The second PCR used primers RU-O-26265 and RU-O-26264 to amplify the RT region of HBV Pol using 50 µl volume and the following cycling conditions: (i) denaturation at 95 °C for 1 min (one cycle); (ii) PCR at 95 °C for 15 s, 62°C for 30 s, and 68 °C for 30 s (12 cycles) (iii) 68 °C for 2 min and 10 °C to hold. See table S3 for primer sequences. DNA from the second PCR reaction was submitted to the CCIB DNA Core Facility, Massachusetts General Hospital (Cambridge, MA, USA) for high throughput amplicon sequencing. Briefly, samples were sheared, ligated to Illumina compatible adapters with unique barcodes, and sequenced from both ends on the MiSeq platform with 2×150 run parameters.

#### DNA sequencing data analysis

Amplicon sequences were mapped to the reference sequence using tophat2 with a maximum of two mismatches allowed. Mapped sequences were read using the RSamtools package (*19*) and split into codons. Codons having one or more base with quality score < 13 were ignored. For paired sample sets (one sample generated in the absence of LAM and one generated in the presence of 400 μM LAM), we performed one-sided Fisher’s exact tests for each possible substitution to obtain p-values for enrichment in the presence of LAM. P-values for substitutions from multiple, independent paired sample sets were combined using weighted z-method (*20*), applied using the metap package (*21*). Source code for variant analysis can be obtained at github.com/ashleyacevedo/HBVseq_sub_enrichment. Sequences are deposited in the Sequence Read Archive, accession number XXX.

## Supplementary figures

**Fig. S1.**
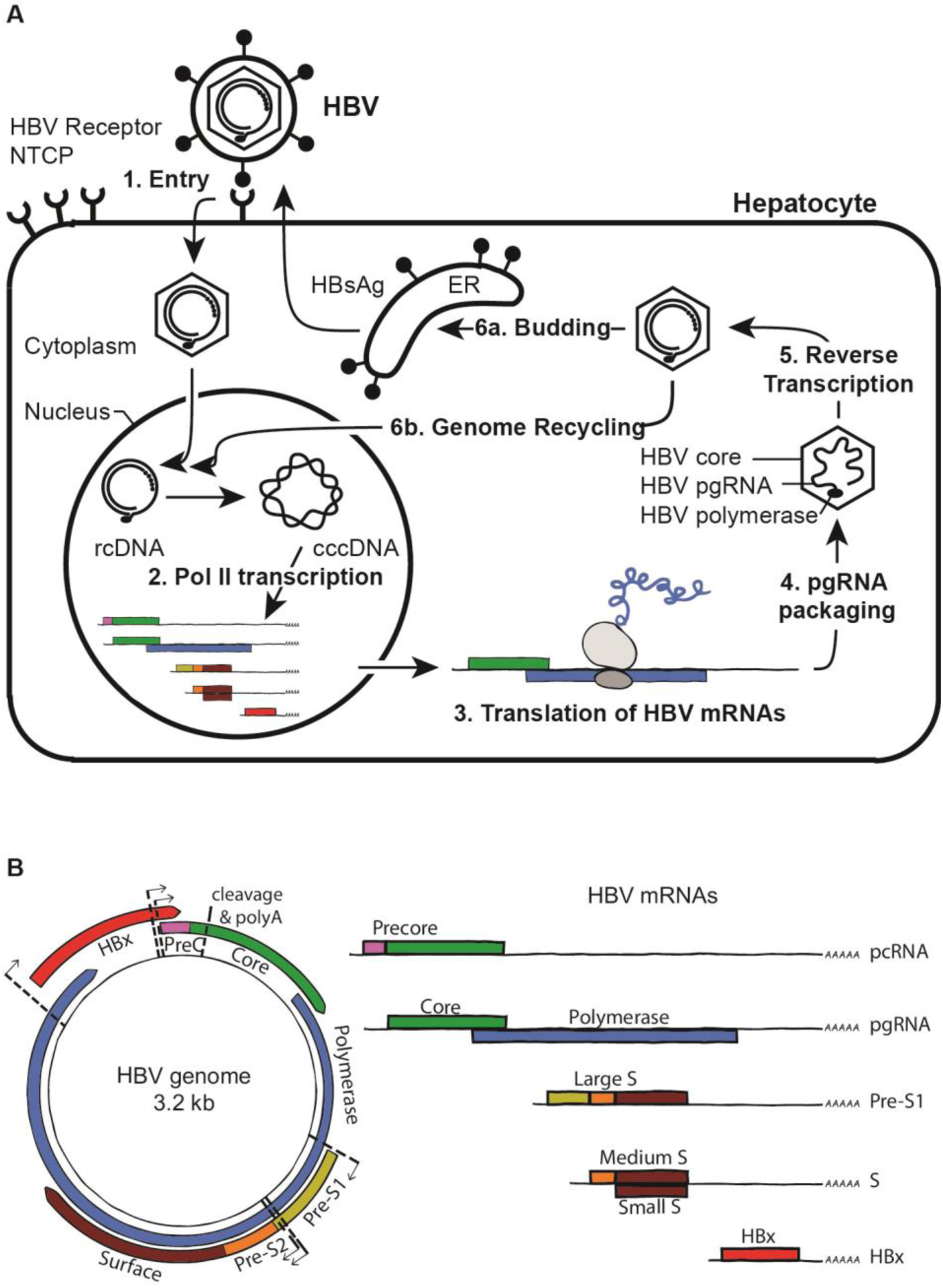
HBV lifecycle and mRNA transcripts. (**A**) HBV lifecycle. HBV (1) enters hepatocytes via heparin sulfate proteoglycans (HSPGs) and sodium taurocholate cotransporting polypeptide (NTCP). In its encapsidated form, the genome is partially double-stranded, and upon entry to the nucleus, is converted from relaxed circular (rc) DNA to covalently closed circular (ccc) DNA. cccDNA is (2) transcribed by cellular Pol II to produce all viral mRNAs, which are then exported to the cytoplasm and (3) translated into viral proteins. Of these mRNAs, pgRNA, serves as the template for reverse transcription and also encodes the core and polymerase proteins. The encapsidated pgRNA (4) is reverse transcribed (5) by HBV polymerase and can either (6a) associate with the viral surface antigens, HBsAg, and be exported from the cell, or (6b) “recycle” to the nucleus and replenish cccDNA. (**B**) HBV genome and mRNA transcripts. Left, the highly compact, 3.2 kb double stranded HBV DNA genome encodes multiple overlapping open reading frames (ORFs). Colored boxes indicate ORFs and arrows indicate Pol II transcriptional start sites. Right, the five HBV mRNAs. All transcripts terminate at the same cleavage and polyA addition signal.

**Fig. S2.**
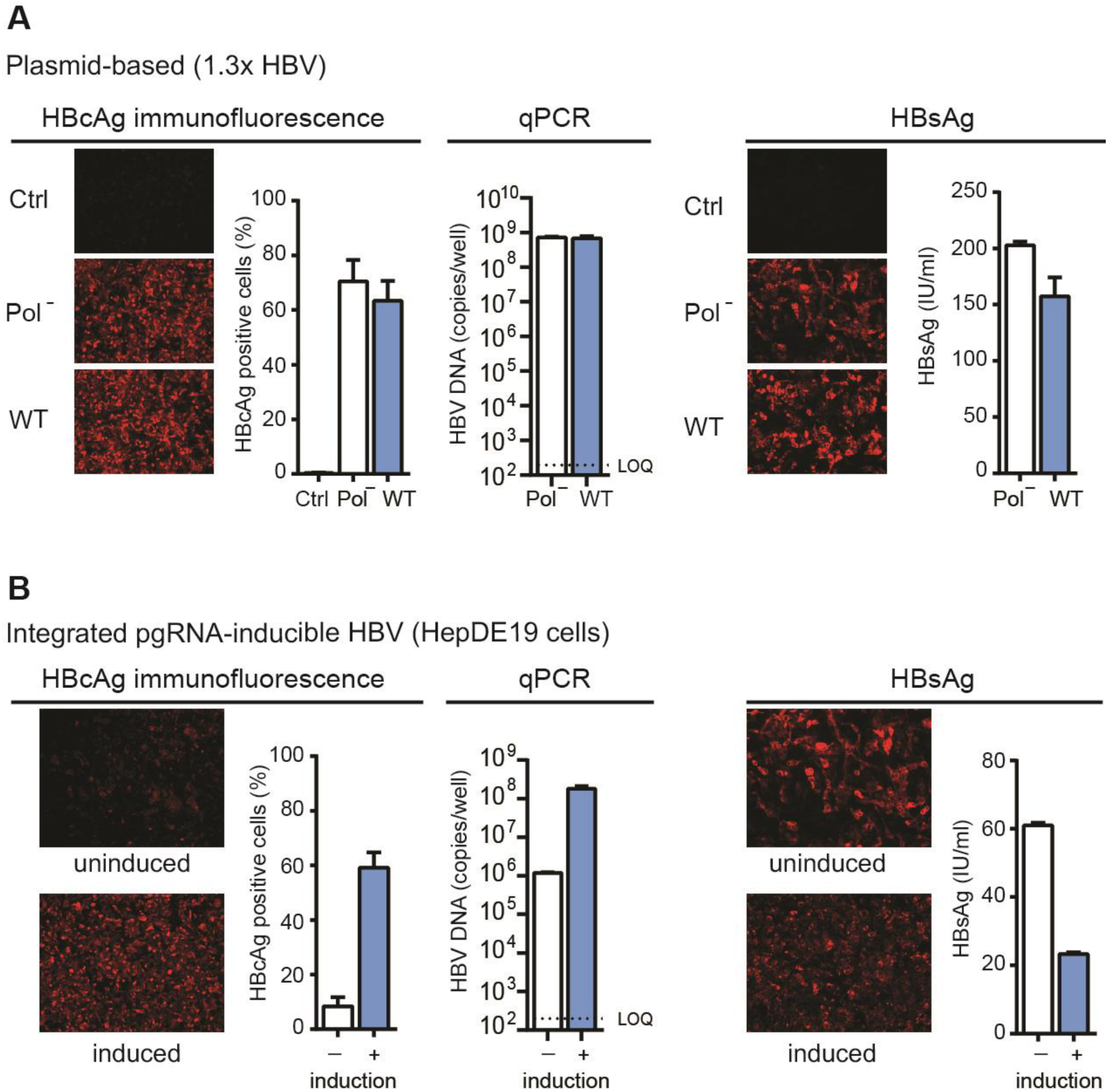
Existing cell culture systems to study HBV. (**A**) A plasmid-based method to initiate HBV replication in cell culture. HepG2 cells were transfected with a plasmid containing over-length (1.3x) HBV genotype A. Anti-HBcAg immunofluorescence indicates that approximately 70-90% of transfected cells stain positive for HBcAg. qPCR detects high HBV DNA copies; however, the signal emanates largely from the transfected plasmid since both WT and Pol^--^ (YMHD) sequences yield similar DNA copy number. Likewise, HBsAg detected in cells by immunofluorescence or in supernatants by CLIA is produced from the transfected plasmid. (**B**) HepDE19 cell line with inducible pgRNA produced from an integrated copy of genotype D HBV. Cells were cultured with (uninduced) and without (induced) tetracycline for five days then either fixed and stained or harvested for DNA extraction. HBcAg immunofluorescence indicates that pgRNA induction increases the percentage of HBcAg-positive cells and qPCR shows a greater than 100-fold increase in HBV DNA copies. (The ∼10^6^ copies of HBV DNA detected in the uninduced condition is likely from HBV DNA integrated in the cellular genome.) Despite clear differences in HBcAg and HBV DNA with and without induction, the HBsAg protein is an unreliable measure of HBV replication since the majority of signal originates from the integrated HBV genome and is independent of replication.

**Fig. S3.**
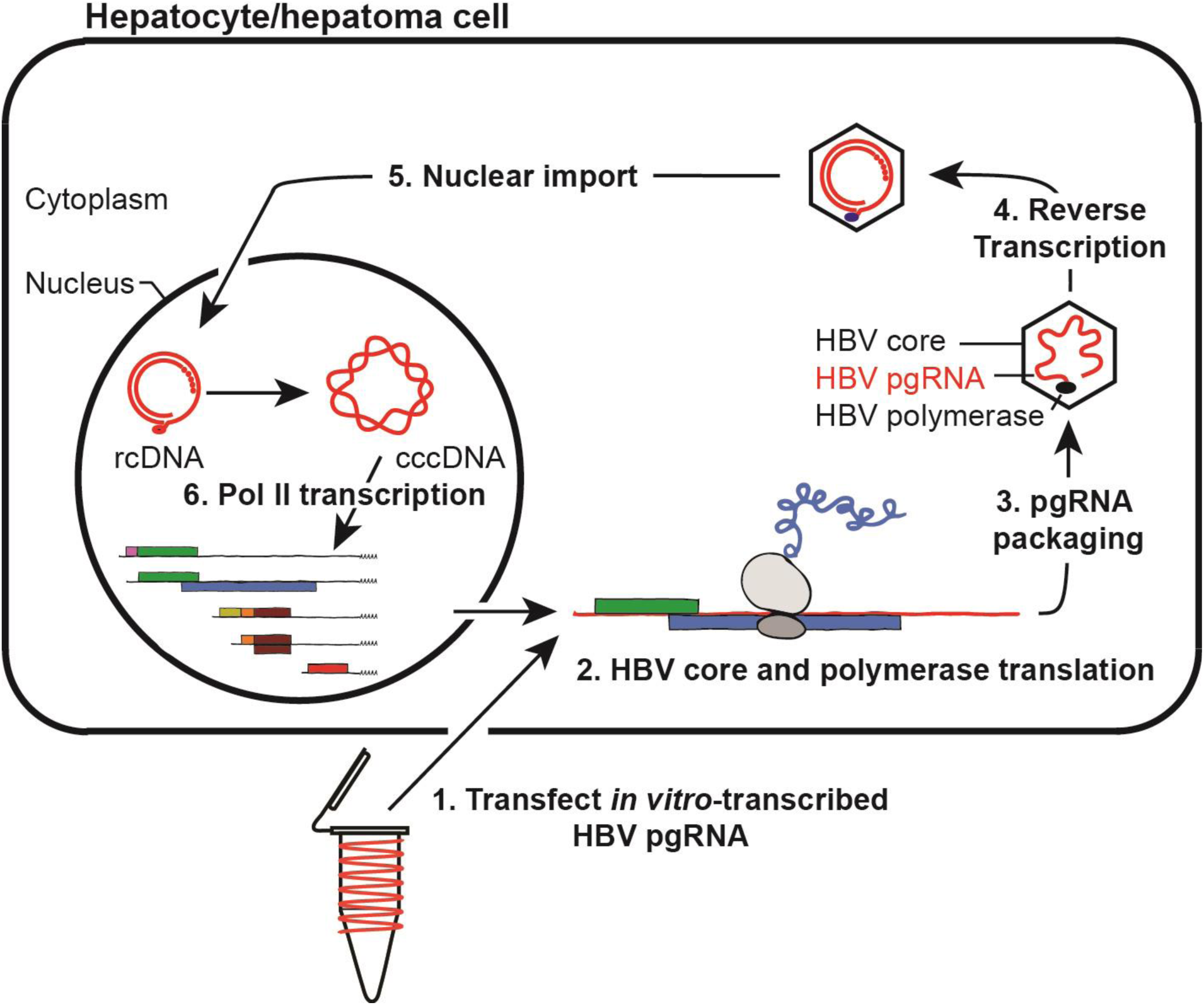
Strategy to initiate HBV infection with pgRNA. *In vitro*-transcribed pgRNA (1) transfected into hepatocytes or hepatoma cells is (2) translated to produce HBV core and polymerase proteins. Together, these two viral proteins (3) encapsidate and (4) reverse transcribe pgRNA into relaxed circular dsDNA. The intracellular virion is then (5) transported to the nucleus where it is converted to cccDNA and (6) transcribed by Pol II to produce all HBV mRNAs.

**Fig. S4.**
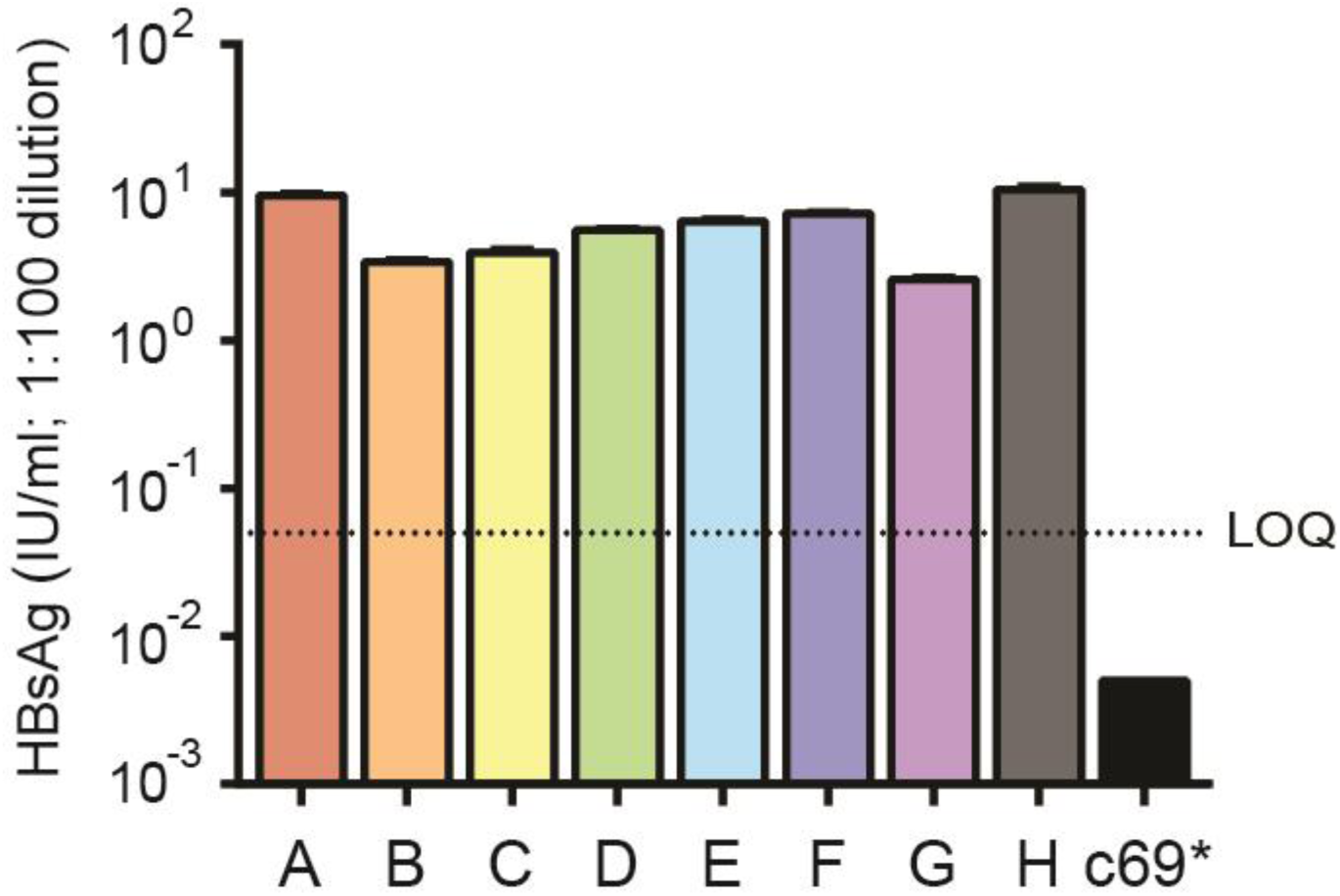
CLIA detects HBsAg from plasmids encoding HBV genotypes A-H. CLIA detects HBsAg from HBV genotypes A-H four days after transfecting cells with the plasmid templates used for *in vitro-*transcription. The results demonstrate that HBsAgs from each genotype are detectable by the CLIA kit and suggest that major differences in HBsAg levels (e.g. genotype G) reflect authentic genotype-specific biology. C69* was included as a negative control. This plasmid (genotype A) encodes a stop codon in the HBs open reading frame at position 69 and therefore does not produce HBsAg. Supernatants were diluted 1:100 prior to detection to measure HBsAg in the same range as the experiment shown in **Fig. 2C**. LOQ, limit of quantification.

**Fig. S5.**
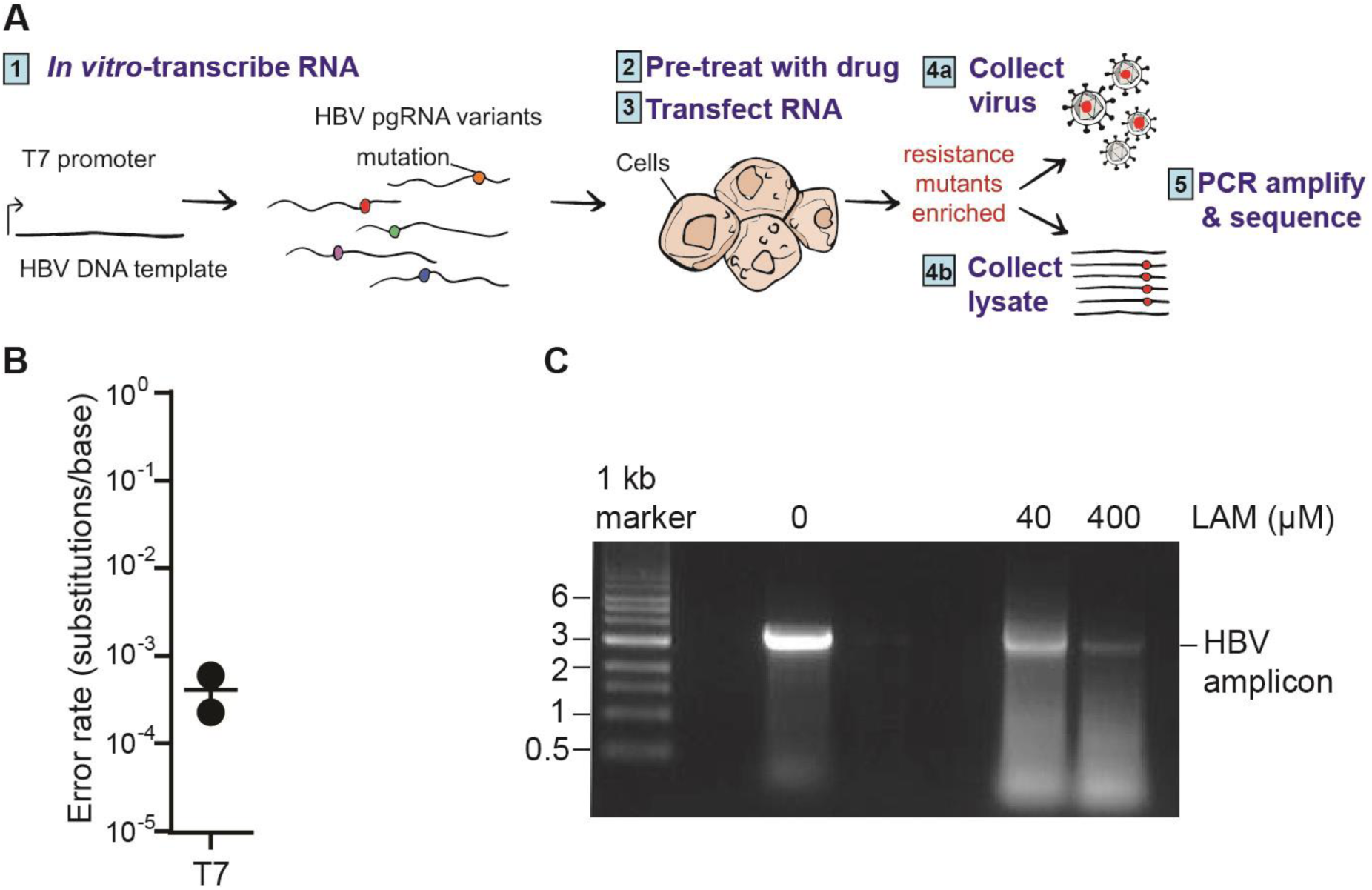
Selecting drug-resistant viral variants from *in vitro*-transcribed HBV pgRNA. **A**) Schematic describes a method to select and enrich mutations that confer resistance to HBV antivirals. From left, T7 bacteriophage RNA polymerase generates a diverse population of HBV pgRNA variants. These RNAs, transfected into cells in the presence of anti-HBV drugs, are subject to selection. HBV DNA, either from cell lysates, in secreted virions, or both, are collected, amplified by PCR, and sequenced with next generation sequencing to identify rare drug-resistant variants. Depending on the selective pressure, reverse-transcribed DNA in the cell, encapsidated DNA in the supernatant, or both, can be PCR amplified and sequenced to enrich for HBV variants. (**B**) Error rate of T7 bacteriophage RNA polymerase determined by CirSeq using *in vitro*-transcribed HBV pgRNA. (**C**) 1% agarose gel of HBV DNA amplified two days post transfection from supernatants of cells untreated or treated with 40 or 400 μM LAM.

**Table S1.** HBV genotypes.

| Genotype | GenBank Accession No. |
| --- | --- |
| A | MN172185 |
| B | MN172186 |
| C | MN172187 |
| D | MN172188 |
| E | MN172189 |
| F | MN172190 |
| G | MN172191 |
| H | MN172192 |

**Table S2.** Dose response of reverse-transcriptase inhibitors.

| log(inhibitor) vs. response<br>(three parameters) | ETV | TDF | LAM |
| --- | --- | --- | --- |
| Best-fit values |  |  |  |
| Bottom | 3.56 | 0.1294 | 0.228 |
| Top | 92.93 | 98.45 | 98.11 |
| LogIC50 | 0.7711 | 1.236 | 3.373 |
| IC50 (nM) | 5.903 | 17.22 | 2363 |
| Span | 89.37 | 98.32 | 97.89 |
| Std. Error |  |  |  |
| Bottom | 4.284 | 4.632 | 2.769 |
| Top | 4.564 | 3.939 | 1.879 |
| LogIC50 | 0.1487 | 0.1309 | 0.0695 |
| Span | 5.924 | 5.75 | 3.17 |
| 95% Confidence Intervals |  |  |  |
| Bottom | -6.924 to 14.04 | -11.20 to 11.46 | -6.548 to 7.004 |
| Top | 81.76 to 104.1 | 88.81 to 108.1 | 93.52 to 102.7 |
| LogIC50 | 0.4072 to 1.135 | 0.9157 to 1.556 | 3.203 to 3.544 |
| IC50 | 2.554 to 13.65 | 8.236 to 35.99 | 1597 to 3496 |
| Span | 74.87 to 103.9 | 84.25 to 112.4 | 90.13 to 105.6 |
| Goodness of Fit |  |  |  |
| Degrees of Freedom | 6 | 6 | 6 |
| R square | 0.9743 | 0.9801 | 0.9941 |
| Absolute Sum of Squares | 341.8 | 314.9 | 84.76 |
| Sy.x | 7.547 | 7.244 | 3.759 |
| Number of points |  |  |  |
| Analyzed | 9 | 9 | 9 |

**Table S3.**
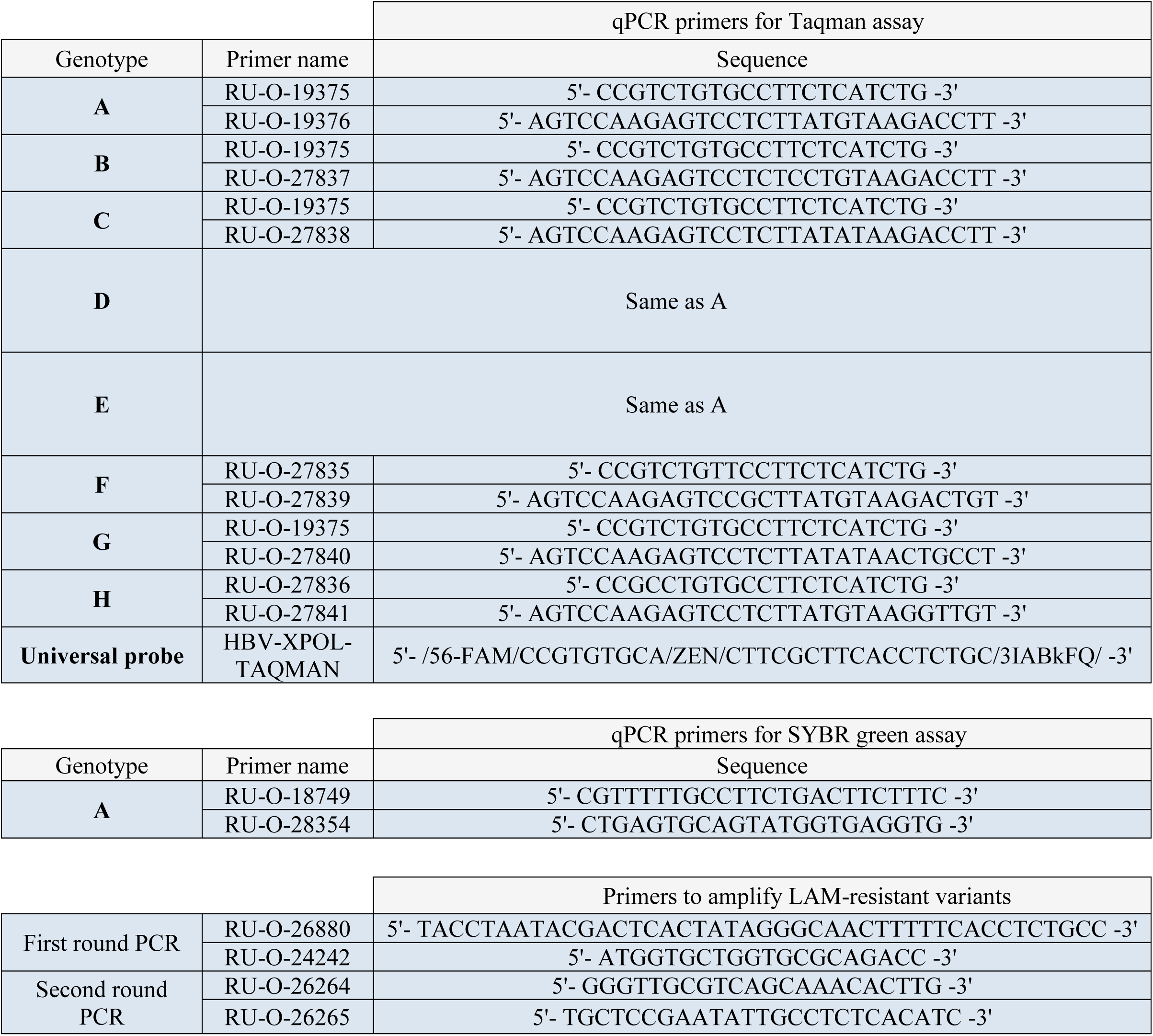
Primer sequences.

